# Decidual cell differentiation is evolutionarily derived from fibroblast activation

**DOI:** 10.1101/2020.12.18.423527

**Authors:** Longjun Wu, Daniel J. Stadtmauer, Jamie Maziarz, Günter P. Wagner

**Author notes:** Günter P. Wagner.

## Abstract

What the molecular mechanisms underlying the evolutionary origin of novel cell types are is a major unresolved question in biology. The uterine decidual cell is a novel cell type of placental mammals which serves as the interface between maternal and fetal tissues during pregnancy. In this paper, we investigate two models for the nature of the differentiation of decidual cells: first, that it represents a mesenchymal-epithelial transition (MET), and second, that it evolved from wound-induced fibroblast activation (WIFA). Immunocytochemistry and RNA-seq analysis of decidualizing human endometrial fibroblasts cast doubt on the MET hypothesis and instead demonstrate a similarity between decidualization and fibroblast activation, including a central role for TGFB1. Through single-cell RNA-seq, we found a transient myofibroblast-like cell population in the *in vitro* differentiation trajectory of human decidual cells and found that these cells represent a pre-decidual state approaching the inferred transcriptomic transition to decidual cells. We propose an evolutionary developmental model wherein the decidual cell is a novel cell type not equivalent to the myofibroblast, but the process of decidual differentiation itself evolved as an endometrial-specific modification to fibroblast activation in response to the wound caused by embryo implantation.

## Introduction

The origin of novel cell types is a major mode for the origin of evolutionary novelties and for the evolution of organismal complexity (Valentine et al., 1994; Arendt, 2008; Wagner et al., 2014; Arendt et al., 2016). The uterine decidual cell (DSC) is a cell type unique to eutherian (Placental) mammals and is critical for the establishment and maintenance of pregnancy in these species (Mess and Carter, 2006; McGowen et al., 2014). DSCs form in response to embryo implantation, or spontaneously during the menstrual cycle in species with menstruation (Gellersen and Brosens, 2014), a derived feature of a minority of eutherian species (Critchley et al., 2020). The evolutionary origin of the DSC along the stem lineage of eutherian mammals was an important step to establish extended pregnancy in eutherian mammals (Wagner et al., 2014; Chavan et al., 2016; Chavan et al., 2020). Decidualization, the process through which endometrial stromal fibroblasts (ESFs) differentiate into DSCs, is a novel cellular response to embryo attachment that does not exist in non-eutherian mammals such as opossum (Kin et al., 2014; Erkenbrack et al., 2018). In this paper we investigate the evolutionary roots of the decidualization process and provide evidence that this process is homologous to, i.e. derived from, the process of WIFA, Wound-Induced Fibroblast Activation.

In previous work, our lab has provided evidence suggesting that the decidualization pathway is derived from an ancestral cellular stress response: ESFs of the opossum, a marsupial whose reproduction represents the inferred ancestral state for viviparous mammals, do not decidualize (Kin et al., 2014), but respond to decidualization signals by activating a cellular stress response (Erkenbrack et al., 2018). Here we address the question of what the biological nature and context of that stress response was.

Resident fibroblasts of mucosal tissues become activated in response to injury, and convert to an activated state called the myofibroblast (Hinz et al., 2007). Myofibroblasts function in wound closure and are marked by the upregulation of proteins such as vimentin (VIM) and alpha-smooth muscle actin (a-SMA) (Darby et al., 2014). Embryo implantation at the beginning of pregnancy causes a local lesion of tissue integrity, i.e. a wound, as the fetus destroys the uterine epithelium and burrows into the stroma (Carson et al., 2000). This traumatic process is the ancestral state for eutherian mammals, as their most recent common ancestor has been inferred to have had hemochorial (invasive) placentation, while non-invasive forms of pregnancy, as observed in hoofed animals and their relatives, evolved later (Wildman et al., 2006; Mess and Carter, 2006; Elliot and Crespi, 2009). Because fibroblasts in all mucosal tissues are activated under similar conditions, it follows that early in the evolution of embryo attachment and implantation, resident fibroblasts of the endometrium would have become activated by the wound caused by embryo attachment. We hypothesize that this routinely-expected stress from tissue injury is the ancestral cell state from which the DSC evolved. We argue that the original function of the DSC was to counteract the inflammation caused by embryo attachment and implantation (Griffith et al., 2017; Chavan et al., 2017; Chavan et al., 2020), an example of stress-induced evolutionary innovation (Wagner et al. 2019).

In this study, we present evidence that decidualization involves induction of some of the characteristic WIFA genes. In addition, we report that transforming growth factor β1 (TGFB1) signaling is involved in decidualization: This is a significant observation because TGFB1 is secreted by epithelial cells upon tissue injury and is a major inducer of fibroblast activation (McCartney-Francis and Wahl, 1994; O’Kane and Ferguson, 1997; Singer and Clark, 1999; Wang et al., 2006). Finally, a single-cell analysis of *in vitro* decidualized cells reveals a transient population of myofibroblast-like cells. These results support the conclusion that WIFA is the developmental as well as evolutionary precursor of the decidualization process. The WIFA derivation model predicts evolutionary links and mechanistic parallels between decidualization, wound healing and cancer malignancy (Kshitiz et al., 2019).

## Results and Discussion

### Decidualization Is Not a Mesenchymal-Epithelial Transition

Our hypothesis that decidualization is derived from WIFA contradicts an alternative hypothesis, namely that decidualization is a mesenchymal-epithelial transition (MET) (Zhang et al., 2013; Pan-Castillo et al., 2018). This idea was inspired by the cell shape changes during decidualization: ESF have fusiform (spindle-shaped) cell bodies typical of mesenchymal cells whereas DSCs are more rounded or “epithelioid”, and express epithelial markers like collagen IV and other basal membrane components (Wewer et al., 1985). Experimental reports of downregulation of genes such as *SNAIL* and *VIM* during decidualization (Zhang et al., 2013; Pan-Castillo et al. 2018) have purported to show that decidualization is a MET-like process. Transcription factors *TWIST1* and *SNAIL* are among the most important regulators of MET and its inverse, EMT, where they shift the balance towards the mesenchymal state (De Herreros et al., 2010; Taube et al., 2010).

To re-test the MET model of decidualization, we analyzed transcriptomes from immortalized human ESF cells decidualized with medroxyprogesterone acetate (MPA, a stable progestin) and 8-Bromo-cAMP (a cell membrane-permeable derivative of cyclic AMP) for 3 and 8 days (Rytkönen et al., 2019). These data show that *TWIST1* and *SNAIL* mRNA expression levels were upregulated (Figure 1A; see also Schroeder et al., 2011), rather than downregulated as expected in a MET. These results were further supported by transcriptomic data from our lab on two primary human ESF cell lines (Figure 1B) (Pavličev et al., 2017), and are consistent with the report of *SNAIL* mRNA being upregulated in the decidual zone of the mouse (Ma et al., 2006). VIM is also an important marker for mesenchymal identity (Bindels et al., 2006; Grande and López-Novoa, 2009). However, contrary to the expectation under the MET model that this mesenchymal marker would decrease, immunocytochemistry showed that VIM protein expression was strongly increased after decidualization (Figure 1C). Consistent with this observation, VIM protein has been reported to be highly expressed *in vivo* in decidual cells of the rat (Glasser and Julian, 1986; Korgun et al., 2007) and human (Can et al., 1995). We note that the *VIM* mRNA level did decrease over decidualization, although at all times exceeded 1000 transcripts per million. We conclude that, during decidualization, the mesenchymal character of endometrial stromal cells is not lost, but enhanced, as documented by the increased VIM protein expression and the upregulation of *TWIST1* and *SNAIL*.

**Figure 1.**
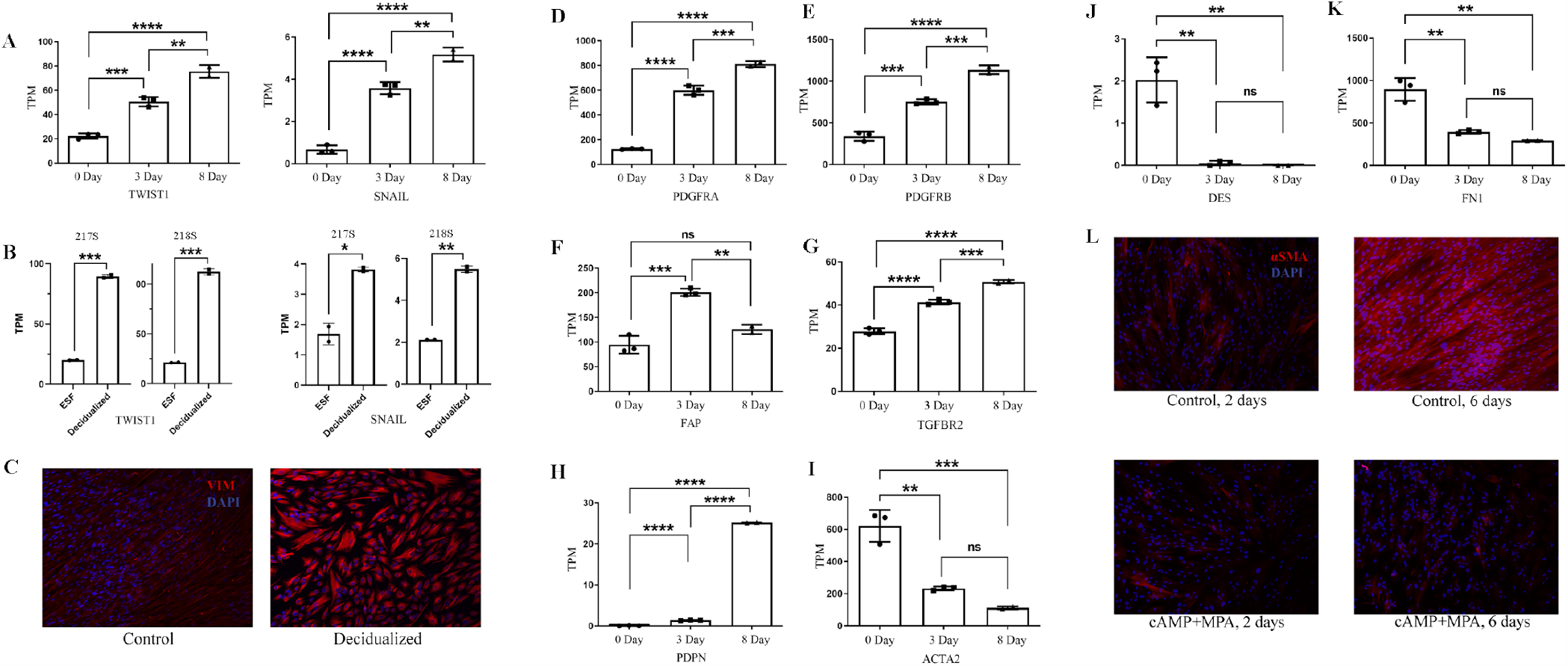
Expression of MET and WIFA pathway components in decidualization. **A-C: MET hallmark genes in decidualization. A:** *TWIST1* and *SNAIL*, transcription factors and markers for fibroblast activation and EMT, are upregulated during *in vitro* decidualization (8-Br-cAMP+MPA, 0, 3, 8 days: Rytkönen et al., 2019). **B:** *TWIST1* and *SNAIL* are upregulated in human primary decidual cell line 217s and 218s during *in vitro* decidualization (Pavličev et al., 2017). **C**: Fibroblast activation and epithelial mesenchymal transition (EMT) marker VIM protein is strongly upregulated during *in vitro* decidualization (8-Br-cAMP+MPA, 8 days). **D-L: Fibroblast activation hallmark genes in decidualization. D-H:** Fibroblast activation regulatory genes *PDGFRA, PDGFRB, FAP, TGFBR2* and *PDPN*, are upregulated during *in vitro* decidualization (8-Br-cAMP+MPA, 0, 3, 8 days: Rytkönen et al., 2019). **I-K:** Phenotypic markers of activated fibroblast cells, *ACTA2, DES* and *FN1* show decreased mRNA expression at later time points of *in vitro* decidualization (8-Br-cAMP+MPA, 0, 3, 8 days: Rytkönen et al., 2019). **L:** Decidual cells (8-Br-cAMP+MPA, 6 days) show low levels of myofibroblast maker a-SMA protein while control cells (6 days) show high levels of a-SMA protein. Early decidual cells (8-Br-cAMP+MPA, 2 days) and control cells (2 days) show low levels of a-SMA protein. Statistical analysis was performed by unpaired t test (C-F) and one-way ANOVA (others). *: p<0.05; **: p<0.01; ***: p<0.001; ****: p<0.0001; ns: not significant.

### Decidualization has Affinities to Fibroblast Activation

Next, we tested whether decidualization is related to WIFA. The WIFA derivation model predicts a transcriptomic signature of fibroblast activation during decidualization. Signaling and regulatory genes of WIFA include the platelet-derived growth factor receptors A and B (PDGFRA and PDGFRB), cell surface receptors essential for wound healing-related fibroblast activation (Floege et al., 2008; Chen et al., 2011; Miyazaki et al., 2019), fibroblast activation protein (FAP) (Tillmanns et al., 2015), and TGF-β receptor II (TGFBR2), a component of TGF-β signaling pathways (Inamoto et al., 2010). Cultured human ESFs increased mRNA expression of all of these genes in response to deciduogenic stimuli (Figure 1D-H), consistent with the WIFA model.

Phenotypic markers for activated fibroblasts also include *ACTA2* (a-SMA), *DES* (desmin) and *FN1* (fibronectin). In accordance with previous findings (Kim et al., 1998; Schwenke et al., 2013), we found a greatly reduced mRNA level during decidualization for all three markers (Figure 1I-K). At the protein level, *in vitro* decidualized cells (8-Br-cAMP+MPA, 6 days) show low levels of a-SMA (Figure 1L). In contrast, control cultures express high levels of a-SMA after 6 days of culture in the medium without MPA or 8-Br-cAMP, but not after 2 days (Figure 1L). This pattern may be explained if the fibroblast activation-like stage of decidualization is transient, as indicated by single-cell analysis of *in vitro* decidualization (see below). The high levels of a-SMA in 6 days of no MPA or 8-Br-cAMP culture compared to 2 days could indicate fibroblast activation by prolonged culture in medium containing only 2% fetal bovine serum, crowding, or by potential active TGFB1 in the medium, which is suppressed under deciduogenic conditions. Next, we turn to a signaling pathway that is critical for wound-induced fibroblast activation, TGFB1.

### TGFB1 Is Involved in Regulating Decidualization

TGFB1 is a cytokine controlling fibroblast activation during wound healing (Brunner and Blakytny, 2004; Oberringer et al., 2008). We reasoned that, if decidualization is derived from fibroblast activation, TGFB1 should also play a role in decidualization, consistent with its role in embryo implantation in various eutherian mammalian species (Singh et al., 2011, Monsivais et al., 2017). We investigated the effects of TGFB1 and pharmacological inhibitors of its receptor on differentiation of human decidual cells *in vitro*. Under prostaglandin E2 (PGE2) and MPA-induced decidualization, TGFB1 protein addition increased mRNA expression of the classic decidual marker *PRL* (Figure 2A), consistent with previous findings (Kim et al., 2005; Chang et al., 2008). In addition, LY2109761, a dual inhibitor of type I and II TGF-β receptors (Melisi et al., 2008) significantly reduced *PRL* mRNA levels (Figure 2B) and reduced the characteristic polygonal morphology of decidual cells (Wynn, 1974) (Figure 2C, D). Hence, TGF-β receptor inhibition weakened decidualization. Taken together, these results support a facilitating role for TGFB1 in human decidualization. The involvement of the TGFB1 signaling pathway in decidualization is consistent with the hypothesis of evolutionary derivation from WIFA.

**Figure 2:**
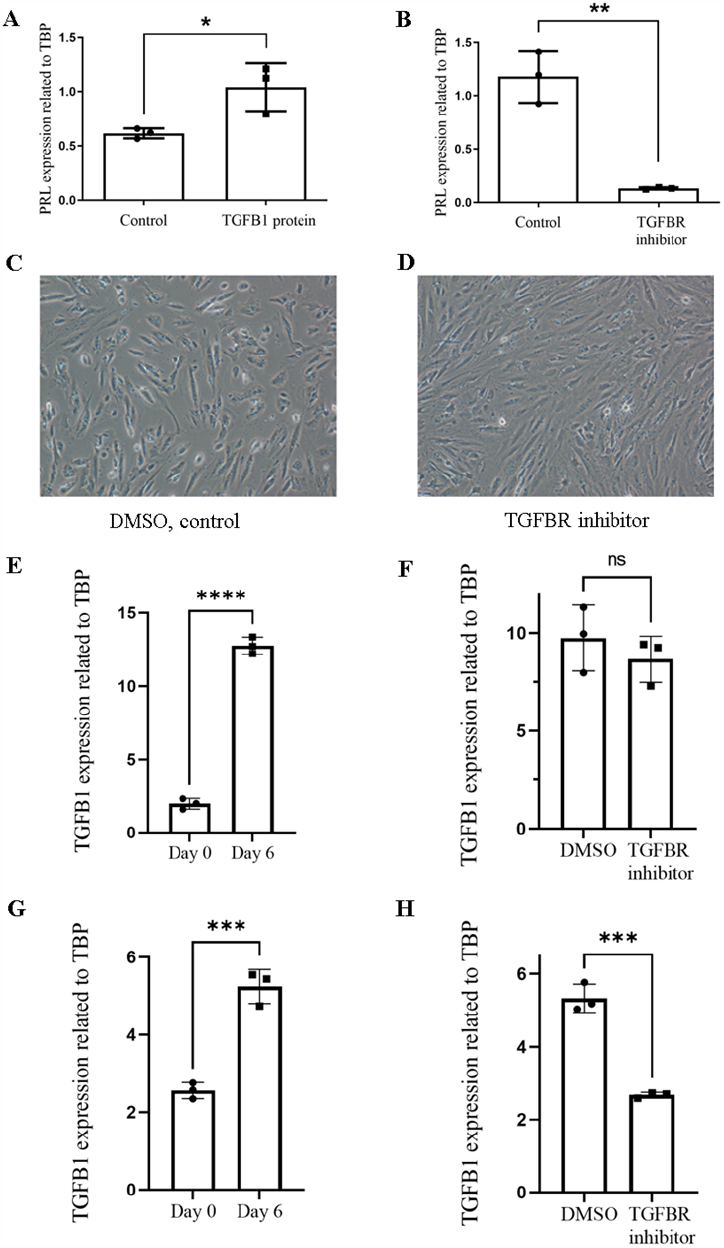
TGFB1 signalling in decidualization. **A-D:** TGFB1 facilitates decidualization. **A)** TGFB1 protein (0.5 ng/mL) increases mRNA expression of decidualization maker *PRL* compared to control (PGE2+MPA, 6 days). **B)** TGF-β receptor inhibitor LY2109761 (5 uM) strongly suppresses mRNA expression of *PRL* (PGE2+MPA, 6 days). **C, D)** Decidual cell polygonal morphology is largely impaired by TGF-β receptor inhibitor (8-Br-cAMP+MPA, 6 days). **E)** In Human ESFs, TGFB1 mRNA expression increased after decidualization (PGE2+MPA). **F)** In Human ESF, TGFB1 expression was not affected by TGF-β receptor inhibitor (PGE2+MPA, 6 days). **G)** In Rabbit ESFs, TGFB1 mRNA expression increased after decidualization (PGE2+MPA). **H)** In Rabbit ESF, TGFB1 mRNA expression was downregulated significantly with TGF-β receptor inhibitor treatment (PGE2+MPA, 6 days). Statistical analysis was performed by unpaired t test, *: p<0.05; **: p<0.01; ***: p<0.001; ****: p<0.0001; ns: not significant.

Given that decidualization increases the expression of *TGFB1* mRNA (Figure 2E) and TGF-β receptor inhibition decreases the expression of decidual markers, we hypothesized that TGFB1 expression itself may be enhanced via an autocrine feedback loop. This seemed plausible given that a positive autocrine feedback loop for TGFB1 has been reported in activated skin fibroblasts during wound healing (Messadi et al., 1994; Moulin, 1995; Blaauboer et al., 2011). However, *TGFB1* mRNA expression in human decidual cells did not show a significant change when treated with the TGF-β receptor inhibitor (Figure 2F), inconsistent with the autocrine feedback of TGBF1 on its own expression.

Next, we asked whether this result is a general feature of the decidualization processes, or whether it is an evolutionarily derived state in humans or other primates. Unlike the vast majority of eutherian mammals, in which decidualization requires the stimulus of an embryo or uterine injury, in humans and a limited number of other menstruating mammals such as some primates, decidualization is independent of the presence of an embryo, or “spontaneous” (Emera et al., 2011). Thus, in humans the increasing expression of TGFB1 in decidualization could be overwhelmingly regulated by hormones or hormone-related pathways over TGFB1 itself.

In order to approach this question, we turned to the ESF cells of the rabbit, a species lacking spontaneous decidualization. We treated primary ESF cells isolated from the rabbit endometrium with PGE2 and MPA. We found that *TGFB1* mRNA expression did increase in rabbit ESFs during treatment with PGE2 and MPA like in human ESFs (Figure 2G). In addition, TGF-β receptor inhibitor was able to block the upregulation of *TGFB1* expression under these conditions (Figure 2H). These results suggest that in the rabbit ESF, increasing TGFB1 expression seems to involve an autocrine feedback loop during PGE2 and MPA treatment, similar to the situation in activated fibroblasts.

These findings, together with the other evidence for an evolutionary relationship between WIFA and decidualization, suggest that the situation in the rabbit ESF could be the ancestral response of mucosal stromal fibroblasts, while the lack of positive feedback in humans could be derived. Clearly, evidence from more species is necessary to resolve the question whether the presence of positive autostimulation is derived or ancestral, but the fact that activated skin fibroblasts have a positive feedback loop of TGFB expression suggests that it is ancestral. On the other hand, in species with spontaneous decidualization, the positive feedback loop may not be necessary: decidualization already happens before implantation rather than in response to embryo attachment and invasion.

### Single-Cell Sequencing Reveals a Myofibroblast-like Cell Population

To further dissect the process of decidualization, we re-analyzed single-cell data from our lab of human ESF decidualized *in vitro* for 2 and 6 days (Stadtmauer and Wagner, 2020). Dimensional reduction and clustering divide the cells into three major clusters: highly proliferative ESF (hpESF), activated ESF (acESF), and DSC, and trajectory analysis infers a single developmental trajectory from ESF to DSC (Figure 3A, B bottom plane). At this point, a population of cells express genes falling into gene ontology (GO) categories such as muscle contraction and actin filament organization (Figure 3C). This expression pattern includes high levels of *ACTA2*, tropomyosin 2 (*TPM2*) and the actin cross-linking protein transgelin (*TAGLN*). This observation suggests that there are myofibroblast-like cells present during the decidualization process; we refer to them as “activated ESFs.”

**Figure 3.**
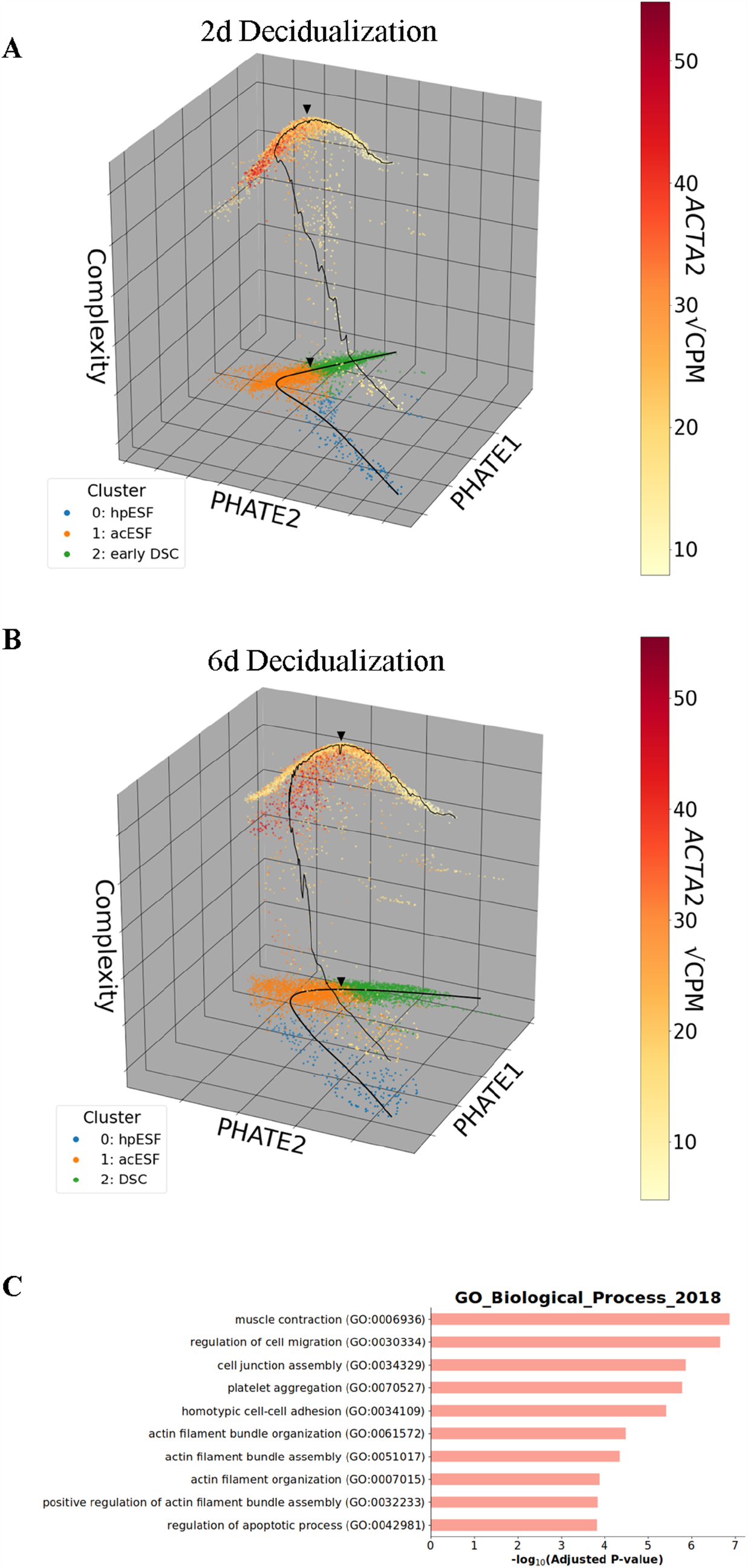
Single-cell RNA sequencing reveals a myofibroblast-like ESF population during Human *in vitro* decidualization. **A, B)** Reduced-dimensionality embeddings for cells 2 days **(A)** and 6 days **(B)** after PGE_2_+MPA decidualization treatment. Cells are plotted in the lower PHATE1-PHATE2 plane, colored by k-means cluster (k = 3), and overlaid with an inferred developmental trajectory. Above, each point is plotted again at a height corresponding to the cell’s intrinsic transcriptomic dimensionality and colored according to expression of the myofibroblast marker *ACTA2* (MAGIC-denoised square-root counts per million). Thus, each cell is represented by a pair of points directly above and below each other. A region of myofibroblast-like *ACTA2+* (red) cells is identified on the ascending side of a complexity peak. The point of highest transcriptomic complexity and its corresponding point in the lower PHATE1-PHATE2 plane are indicated by arrowheads. **C)** Output from gene ontology analysis of the myofibroblast-like cell cluster from the 6-day sample, ranked by Enrichr P-value adjusted for multiple comparisons.

This activated ESF population observed *in vitro* corroborates two single-cell sequencing studies of the first-trimester human fetal-maternal interface which also reported myofibroblast-like cells (Suryawanshi et al., 2018; Vento-Tormo et al., 2018). These myofibroblast-like cells are characterized by genes such as *ACTA2* and *TAGLN*. The presence of such cells *in vivo* suggests that our identification of a myofibroblast-like ESF cell population is not an *in vitro* artifact; on the contrary, our *in vitro* finding suggests that the *in vivo* finding of myofibroblast-like cells may reflect part of the decidualization process rather than activated tissue fibroblasts from other parts of the uterus.

As these studies of the first-trimester human fetal-maternal interface were *in vivo* observational data rather than controlled differentiation studies, the developmental trajectories between these myofibroblast-like ESFs and DSCs could not be clearly ascertained. To assess whether myofibroblast-like ESFs are a distinct population or transitory state, we took a closer look at our single cell experimental decidualization data. Intrinsic dimension (Kerstin, 2016) of a cell transcriptome is a measure of transcriptomic complexity reflecting the number of genes simultaneously expressed. A high level of transcriptomic complexity has been related to cells undergoing transcriptomic reprogramming to assume a new cell type identity (Brunskill et al., 2014), as they will transiently co-express two transcriptional programs. Based on this insight, and previous work using intrinsic dimensionality to identify branch points (Moon et al., 2019), we visualized single-cell transcriptomes of decidualizing human ESFs as a landscape — a two-dimensional PHATE embedding (Moon et al., 2019) and in the third dimension a measure of intrinsic dimensionality (Kerstin, 2016; Moon et al., 2019). The resulting landscape (Figure 3A, B) contained a single point of high transcriptomic complexity (indicated by arrowheads), corresponding to the ESF-DSC developmental transition (Stadtmauer and Wagner, 2020). The myofibroblast-like cells fell squarely into the ESF side of this transition point, as visualized by the marker *ACTA2* (highlighted in red) characterizing the ascent of the complexity hill but not its descent towards DSC (Figure 3A, B). The location of these cells suggests that myofibroblast-like ESFs represent an activated ESF state transitional to DSC, rather than an alternative cell type fate. We interpret this as consistent with our model that the process of decidualization is an evolutionary modification of WIFA.

The single-cell transcriptomic data suggest that decidualization is a two-step process. First, ESFs assume an activated pre-decidual state, characterized by a myofibroblast-like gene expression profile, which is followed by differentiation into the actual DSC. The latter process departs from this activated, myofibroblast-like state, rather than from the quiescent ESF. Such a scenario would also explain the partial affinity between the DSC and WIFA, which are similar at the signaling and transcription factor level but divergent in their functional effector phenotype. Whether this two-phase process corresponds to the idea that decidualization is a biphasic process with an early inflammatory and a later anti-inflammatory phenotype (Salker et al., 2012, Rytkönen et al., 2019) will require more detailed investigation. The fact that myofibroblast-like cells have been identified in the *in vivo* implantation site (Suryawanshi et al., 2018; Vento-Tormo et al., 2018) suggests that the second phase of the process (transition from the activated pre-decidual state to DSC) could be reversible: a dynamical equilibrium like a chemical reaction.

Generation of a myofibroblast-like ESF cell state *in vitro*, distinct from DSC, has also been achieved through exposure of ESFs to platelets, proposed to act through and dependent upon TGFB1: excess production of these myofibroblast-like ESF was proposed to drive a constitutive wound healing phenotype in endometriotic lesions (Zhang et al., 2016). It is possible, therefore, that normal progression from the fibroblast activation-like stage to decidualization and prevention of its persistence (Nancy et al., 2018) is dependent upon proper receipt of uterine-specific regulatory factors. The physiological role of myofibroblast-like pre-decidual cells warrants further study.

### A Model for the Evolution and Nature of the Decidualization Process

We present our evolutionary/developmental model in Figure 4. Based on this model, we suggest that the developmental process of decidualization likely derived from WIFA caused by the implanting embryo. Our single-cell analysis suggests that the decidualization process includes at least two steps: one where the cells assume a myofibroblast-like state, probably homologous to the generic WIFA, and from that cell state the actual differentiation of the mature decidual cell proceeds. Hence, we have the following cell state changes: 

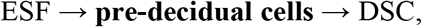

where the first step corresponds to

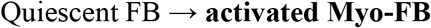

during the wounding response. This model suggests that the pre-decidual cell is homologous to the activated fibroblast, and the DSC is a novel cell type arising on the foundation of activated fibroblast. Based on the phylogenetic distribution of placental types (Wildman et al., 2006; Mess and Carter, 2006; Elliot and Crespi, 2009) we infer that this evolutionary event must have occurred in the stem lineage of eutherian mammals. While some similarities between myofibroblasts and mature DSC have been previously noted (Oliver et al., 1999), this model does not require identifying myofibroblasts as a sister cell type to DSC, something which a previous cell type phylogenetic analysis did not support (Kin et al., 2015). In contrast to generic fibroblast activation, decidualization occurs after estrogen priming and under sustained progesterone stimulation (Gellersen and Brosens, 2014). The latter signal indicates to the endometrium that ovulation has happened and eventual wounding of the uterine epithelium might be caused by an embryo. Indeed, wounding in the uterus in the presence of high progesterone levels is sufficient to cause decidualization, as demonstrated in the so-called deciduoma model of rodent decidualization (Krehbiel, 1937; Finn, 1986). Our model explains at the one hand the numerous mechanistic similarities between decidualization and wound healing, because of their homology, but on the other hand also accommodates the fact that mature decidual cells display a distinct phenotype at maturity adapted to their role in embryo implantation. Among them are the active suppression of genes dedicated to the actual wound healing process (Nancy et al., 2018). Given the possible linkage of wound healing to both decidualization and to cancer malignancy (Dvorak, 1986, 2015), we note that suppression of wound healing may be relevant for cancer progression.

**Figure 4.**
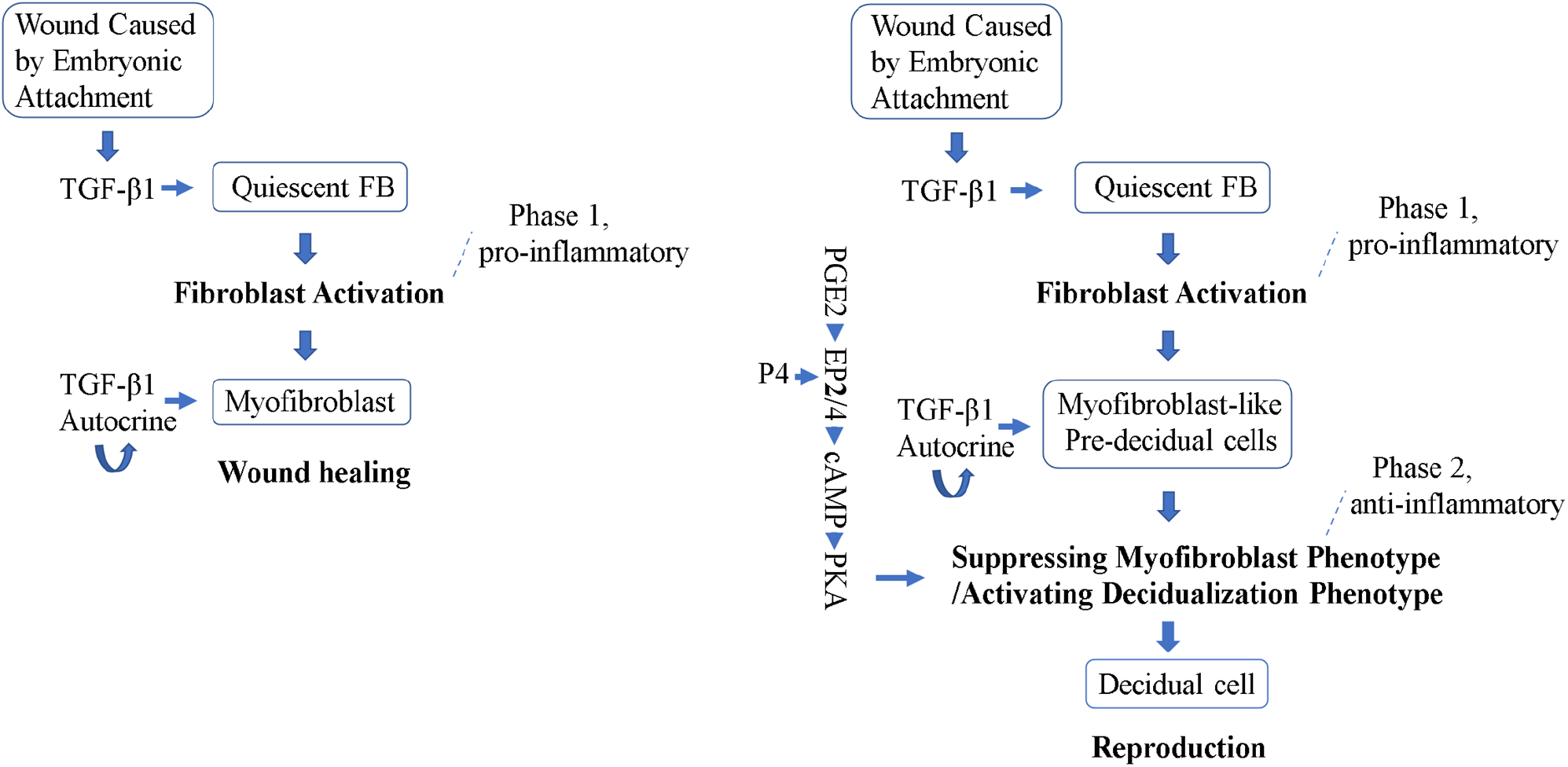
A developmental and evolutionary model for decidualization. Two stages of the evolution of the embryonic attachment reaction are shown. In the first **(left)** the attachment leads to a generic fibroblast activation and wound healing, in the second **(right)** the attachment reaction is transformed into decidualization. Hormone regulations required for the second phase. P4: progesterone, PGE2: Prostaglandin E_2_, E2: Estradiol, EP2/4: Prostaglandin E2 receptors 2/4.

This model of novel cell type evolution is distinctive from other models by emphasizing intermediate processes: transient cell states which could be homologous to pre-existing cellular processes. Since cellular processes are responses to stimuli (an environment under which novel cell types could thrive), we hypothesize that novel cell types may often build upon the foundation of transient cell states. Both activation, as in WIFA, and differentiation are types of cell state transitions differing primarily in the stability of the assumed state (Pope and Medzhitov, 2018), and the origin of differentiation processes from activation processes may be a repeated trend in evolution. For instance, there is growing evidence that cellular stress responses can be the seed for the evolutionary origin of novel cell types (e.g. Wagner et al., 2019; König and Nedelcu, 2020). Furthermore, under this model, it seems that the transient cell state does not necessarily need to be stress-induced: any cellular process induced by internal or external signaling may do.

Dynamic transient cell states like this could have been missed in studies based on bulk cell populations such as western blots, qPCR and bulk RNA-seq analysis. The present study highlights the important insights that become possible from employing single-cell analysis to capture these transient states. In sum, a single-cell level of analysis may open up a new direction to understand the evolutionary origin of novel cell types.

## Acknowledgements

We thank the members of the Wagner lab for support and critical discussions about the subject of this paper, as well as the staff of the Yale Center for Genome Analysis for their excellent work in Next-Gen Sequencing. This work was supported by the John Templeton Foundation research grant #61329 to GPW. The work does not represent the views of the John Templeton Foundation. Additional support for the work related to this study includes NCI grant U54-CA209992.

## Materials and Methods

### Cell culture

Immortalized human ESFs (ATCC CRL-4003) were cultured in growth media containing 1x antibiotic-antimycotic (15240, Gibco; contains penicillin, streptomycin, and Amphotericin B), 15.56 g/L DMEM/F-12 without phenol red (D2906, Sigma-Aldrich), 1.2 g/L sodium bicarbonate, 10 mL/L sodium pyruvate (11360, Thermo Fisher), and 1 mL/L ITS supplement (354350, VWR) in 10% charcoal-stripped fetal bovine serum (100-119, Gemini).

ESFs from rabbit (*Oryctolagus cuniculus*) were isolated from uterine tissues by enzymatic digestion. The uterus was cut in half longitudinally and then cut into 2-3 mm fragments. These were digested with 0.25% trypsin-EDTA for 35 min at 37°C, followed by dissociation buffer (1 mg/mL collagenase, 1 mg/mL Dispase, 400 mg/mL DNaseI) for 45 minutes at 37°C. Cells were homogenized by passing through a 22-gauge needle and syringe followed by a 40 mm nylon mesh filter to remove remaining clumps. A single-cell suspension was obtained from the lysate, transferred to fresh growth medium, and cultured in T25 flasks. To facilitate enrichment of fibroblasts over epithelial cells, media was exchanged after fifteen minutes in order to remove less adherent cells (differential attachment). Cells were grown to confluency and sub-passaged by scraping the cells off of the surface and splitting into 2 T25 flasks, and one additional round of differential attachment was performed. All endometrial stromal fibroblasts were cultured in growth media (as above), with no antibiotic-antimycotic. To validate our fibroblast cell line, we used immunocytochemistry to detect proteins that mark mesenchymal (vimentin+) rather than epithelial (cytokertain+) cells cells. Primary antibody solutions used as follows: 1:200 rabbit anti-cytokeratin (ab9377, Abcam); 1:200 mouse anti-vimentin (sc-6260, Santa Cruz).

### Decidualization treatment

Decidualization was effected with medium containing antibiotic-antimycotic (15240, Gibco), 15.56 g/L DMEM/F-12 (D8900, Sigma-Aldrich), and 1.2 g/L sodium bicarbonate in 2% fetal bovine serum (100-106, Gemini). Control wells were grown in medium alone while experimental wells additionally received 0.5 mM 8-Br-cAMP (B7880, Sigma-Aldrich) and 1 μM MPA (M1629, Sigma-Aldrich), or 1 μM PGE2 (HY-101952, MedChemExpress) and 1μM MPA. Media was replenished every two days.

### Immunocytochemistry

Human ESFs were grown in an 8-well chamber slide (12-565-18, Fisher) to 70-80% confluency before treatment and decidualized for the duration indicated. Cells were fixed with 4% paraformaldehyde in phosphate-buffered saline (PBS) for 15 mins at room temperature after treatment. Cells were then washed three times with PBS. Permeabilization was performed with 0.25% Triton X-100 in PBS for 10 mins at room temperature, followed by three more washes with PBS. For blocking, cells were incubated with 1% BSA in PBST (PBS with 0.1% Tween 20) for 30 mins at room temperature. Primary antibody incubation (1:100 mouse-anti-Vim: sc6260, Santa Cruz or 1:100 mouse-anti-aSMA: ab7817, Abcam) was performed with the aforementioned blocking solution at 4°C overnight. Cells were then washed three times with PBS. Secondary antibody incubation (1:100 Alexa Fluor 555 goat anti-mouse: A21422, Thermo Fisher) was performed with blocking solution at room temperature for 1.5 hours. Cells were washed with PBS three times before staining with DAPI (10236276001, Roche) for 1 min. Cells were then washed with PBS three times and mounted with 50% glycerol. Staining results were imaged on a Nikon Eclipse E600 microscope equipped with a Spot Insight camera.

### TGFB1 Treatment

TGFB1 experiments were performed in media containing 1 μM PGE2, 1 μM MPA, 1x antibiotic-antimycotic, 15.56 g/L DMEM/F-12 (D2906) without phenol red, 1.2 g/L sodium bicarbonate, 10 mL/L sodium pyruvate, 1 mL/L ITS supplement, and with no fetal bovine serum to exclude the effects of potential active TGFB1 in fetal bovine serum. Recombinant human TGFB1 protein (ab50036, Abcam) was added to the media of experimental wells at the beginning of the treatment (0.5 ng/mL). Control wells received the same media but no TGFB1 protein. For the TGF-β receptor inhibition experiment, LY2109761 (S2704, Selleckchem) was added into the media at the beginning of the treatment (5 μM). 0.5 μL DMSO was added in the control group. Decidual cell morphology experiment was performed under decidualization treatment.

### Quantitative PCR

Human ESFs were cultured to 70-80% confluency before treatment. After treatment, media was removed and cells were washed two times with PBS and then lysed with Buffer RLT Plus plus 1% beta-mercaptoethanol. RNA was extracted using an RNeasy Plus Micro Kit (74034, Qiagen), and cDNA was constructed with a High-Capacity cDNA Reverse Transcription kit (4368814, Applied Biosystems). qPCR was carried out in triplicate using ThermoFisher TaqMan reagent on StepOne Plus Real Time PCR System 482 (Applied Biosystems). Gene TBP was used as the internal reference gene to normalize the expression of target genes. qPCR primers used are as follow: Human *TBP*: Hs99999910_m1, Human *PRL*: Hs00168730_m1, Human *TGFB1*: Hs99999918_m1, Rabbit *TBP*: Oc05211505_m1, Rabbit *TGFB1*: Oc04176122_u1.

### Single cell transcriptomic analysis

Published sequence data (NCBI PRJNA679324) of human ESF treated with PGE2+MPA daily for 2 or 6 days were aligned and pre-processed using scprep software version 1.0.5 (github.com/KrishnaswamyLab/scprep) following the original study to filter irregularly-sized libraries resulting from dead cells or doublets (Stadtmauer and Wagner 2020). Briefly, thresholding between the 16^th^ and 90^th^ percentile of features per library for 2-day treated samples, between the 9^th^ and 90^th^ percentile of features per library for 6-day treated samples, and below the 93^rd^ percentile of mitochondrial gene content of all libraries resulted in 4077 remaining cells from the 6-day group, and 3202 cells in the 2-day group, from a combined total of 10073. Thresholding to genes only expressed in 5 or more cells left 16496 genes in consideration. Read counts were normalized to square root counts per million.

A two-dimensional embedding was calculated using PHATE (v1.0.5, Moon et al., 2019). K-means clustering was performed with a hyperparameter of 3, and differential gene expression between clusters was calculated using scprep’s differential expression function which performed Welch’s *t*-test between cells within and without each cluster. Gene set enrichment analysis on the markers of the cluster identified as activated ESF was conducted by querying Enrichr (Kuleshov et al., 2016). Intrinsic dimensionality for landscape plotting was calculated following Moon et al. (2019), using a method based on the intrinsicDimension R Package and Kerstin (2016). A developmental trajectory between clusters was drawn using Slingshot (Street et al., 2018) with the starting cluster set to that with lowest dimensionality, and interpolation was used to draw the trajectory in the intrinsic dimensionality axis as well to aid visualization. Expression of *ACTA2* was denoised using MAGIC (v2.0; van Dijk et al., 2018) for clarity of visualization.

## Notes

### Competing Interest Statement

The authors have declared no competing interest.

